# A Novel CD206 Targeting Peptide Inhibits Bleomycin Induced Pulmonary Fibrosis in Mice

**DOI:** 10.1101/2020.07.27.218115

**Authors:** Anghesom Ghebremedhin, Ahmad Bin Salam, Benjamin Adu-Addai, Steve Noonan, Richard Stratton, Md. Shakir Uddin Ahmed, Chandra Khantwal, George R Martin, Huixian Lin, Chris Andrews, Balasubramanyam Karanam, Udo Rudloff, Henry Lopez, Jesse Jaynes, Clayton Yates

## Abstract

Activated M2 polarized macrophages are drivers of pulmonary fibrosis in several clinical scenarios such as Acute Respiratory Disease Syndrome (ARDS) and Idiopathic Pulmonary Fibrosis (IPF), through the production of inflammatory and fibrosis-inducing cytokines. In this study, we investigated the effect of targeting the CD206 receptor with a novel fragment of a Host Defense Peptide (HDP), RP-832c to decrease cytokines that cause fibrosis. RP-832c selectively binds to CD206 on M2 polarized bone marrow derived macrophages (BMDM) *in vitro*, resulting in a time-dependent decrease in CD206 expression, and a transient increase in M1 marker TNFα, which resolves over a 24hr period. To elucidate the antifibrotic effect of RP-832c, we used a murine model of bleomycin (BLM) -induced early-stage pulmonary fibrosis. RP-832c significantly reduced bleomycin-induced fibrosis in a dosage dependent manner, as well as decreased CD206, TGF-β1 and α-SMA expression in mouse lungs. Interestingly we did not observe any changes in the resident alveolar macrophage marker CD170 expression. Similarly, in an established model of lung fibrosis, RP-832c significantly decreased fibrosis in the lung, as well as significantly decreased inflammatory cytokines TNFα, IL-6, IL-10, INF-γ, CXCL1/2, and fibrosis markers TGF-β1 and MMP-13. In comparison with FDA approved drugs, Nintedanib and Pirfenidone, RP-832c exhibited a similar reduction in fibrosis compared to Pirfenidone, and to a greater extent than Nintedanib, with no apparent toxicities observed on body weight or blood chemistry. In summary, RP-832c is a potential agent to mitigate the overactivity of M2 macrophages in pathogenesis several pulmonary fibrotic diseases, including SARS-CoV-2 induced lung fibrosis.

## Introduction

Multiple fibrotic diseases such as acute lung injury (ALI), acute respiratory distress syndrome (ARDS) and idiopathic pulmonary fibrosis (IPF), share many pathophysiological features including a pro-inflammatory stimulus leading to a rapid release of IL-8 and IL-6 by alveolar macrophages that further attract neutrophils causing alveolar and endothelial injury [1], ultimately resulting high mortality rates. Although the majority of IPF patients have a relatively slow progression of disease, there is a subset of rapid progressors that demonstrate upregulation of inflammatory pathways and have an accelerated loss of lung function and shorter survival [2, 3] similar to ALI/ARDS patients. The survival of ARDS patients or rapidly progressing IPF patients is directly attributed to the deposition of dense parenchymal fibrosis and ultimate loss of pulmonary function. This is currently being observed by the world as SARS-CoV-2/COVID-19 infected patients exhibit many of these features that progress rapidly causing the patient to require mechanical ventilation. Thus, there is great need for therapeutics which block the inflammatory features in these conditions.

The fibroproliferative phase of ALI/ARDS has traditionally been regarded as a late event. However, recent studies suggest that ARDS patients display increased collagen turnover within 24 hrs of diagnosis [4], with increased levels of active TGF-β1 in both the bronchoalveolar lavage fluid (BALF) and tissue samples [5]. Macrophages also regulate fibrosis by secreting growth factors and cytokines, including TGF-β1, that recruit and activate fibroblasts and other inflammatory cells [6], which in turn promote collagen-producing myofibroblasts [7, 8]. Macrophages can exhibit a variety of phenotypes with M1 classically activated macrophages classified using canonical markers such as IFN-γ, CD80, and CD86. M2 alternatingly activated macrophages show a high expression of CD206 expression in disease promoting M2 and induce immunosuppression [9, 10]. Several studies have demonstrated that activated CD206 positive M2 macrophages in fibrotic lesions, produce high amounts IL-10, IL-6, TNF-α, and TGF-β1 that enhance collagen synthesis and deposition [11, 12]. These pro-inflammatory and profibrotic cytokines produced by M2-macrophages indirectly inhibit the production of anti-inflammatory cytokines in a negative feedback further promoting fibrosis [13–15].

Host Defense Peptides (HDP) are ubiquitously expressed in many complex organisms and are critical mediators of the innate immune response [16]. Recently several groups have identified fragments of these HDPs (10-15 aa.) have immunomodulatory activity [16, 17]. These 10-15 aa fragments are also found in internal sequences of collagens, complement, and virulence factors, including pathogens such as bacteria and viruses, that induce cellular changes in many immune cell types including leukocytes, and macrophages [16, 18]. Our group reported that HDP, RP-182 and RP-832c, specifically target the CD206 receptor on M2 macrophages, inducing a major conformation change activating a signaling pathway which rapidly induces apoptosis of CD206 positive M2 macrophages as well as repolarization towards the M1 phenotype[18]. While the efficacy of these peptides has been extensively characterized in multiple tumor models, it’s not well characterized in lung fibrosis. The bleomycin model of bleomycin (BLM)-induced inflammation and fibrosis represents an experimental model for IPF as well as ALI/ARDS [19, 20], therefore we used this model to more completely define the role of RP-832c in lung fibrosis.

## Material and Methods

RP-832c, RWKFGGFKWR peptide was synthesized by PolyPeptide Laboratories, San Diego, CA. BMDM cells were isolated and polarized as previously described [18]. Cell viability of RP-832c treated macrophages were determined using the live/dead viability assay as previously described [18]. Human fetal lung fibroblast cell lines MRC5, IMR90, and IMR9 were purchased from the ATCC (Manassas, VA), and were cultured according ATCC protocols.

### Animal Experiments

C57BL/6J mice were obtained from Envigo, USA. All the animal studies were approved by the IACUC (Murigenics, Inc). Mice were challenged with a single 2.5U/kg body weight dose of bleomycin (BLM) intratracheally (IT) and assessed for fibrosis either 3 days or 14 days post BLM. RP-832c was inoculated subcutaneously for either 21 or 14 days. Body weights were measured daily, and the lung weight was measured at the conclusion of each study.

### Histological and Immunostaining Evaluation

Lungs were sectioned and stained with both Hematoxylin/Eosin (H&E) (Sigma Aldrich, USA) and Masson Trichrome (Abcam, USA). Immunostaining was performed with the anti-CD206, CD107, TGF-β1 and α-SMA R&D systems antibodies as previously described [18]. All slides were scanned using a Leica Aperio SC2 scanner and blindly evaluated by a pathologist. Immunofluorescence images represent 10 fields per tissue section and individually analyzed for total area using confocal microscope (Olympus, New York). All quantitative data were normalized to appropriate control images.

### Quantitative Real Time PCR

Quantification of individual mRNA expression was performed by real-time PCR on an ABI 7500 Fast Real Time System (Applied Biosystems, Foster City, CA, USA), using TaqMan probes as previously described [18].

### Statistical Analysis

Data are reported as means± SD. Comparisons between the groups were performed using a paired t-test. All statistical analyses were conducted using GraphPad Prism software version 8. *P* < 0.05 was considered statistically significant.

## Results

### RP-832c targets CD206 positive macrophages

Since RP-182 is well characterized, we sought to further characterize the physical and functional properties of RP-832c. RP-832c is an amphipathic β-sheet of two palindromic pentamers and is predicted to bind phenylalanine residues (821, 831 and 861) in the CRD5 binding region of CD206 (Figure 1A). In silico docking studies predicted RP-832c to have a higher affinity/COB to full length CD206 compared RP-182. Surface plasma resonance (SPR) binding analysis confirmed that RP-832c, binds recombinant CD206 receptor protein rapidly in a dosage dependent manner (Supplemental Figure 1A), with an estimated *K*_D_ of 3.5μM (Figure 1B). To determine specificity of RP-832c for M2 macrophages, we performed live dead assays using bone marrow derived macrophages polarized to either M1 or M2 phenotypes (Supplemental Figure 1B). RP-832c exhibited a dosage dependent inhibition in the cell viability in M2 macrophages, with an IC50 of ~6 μM producing only a minimal cytotoxicity in M1 polarized macrophages (Figure 1C, D); or two different human fetal lung fibroblast cell lines MRC5 and IMR9 up to 100μM (Supplemental Figure 2A, B). RP-832c further decreases CD206 expression on M2 macrophages (Figure 1) resulting in an initial increase in TNF-α, which declines over a 24hr period (Figure 1G). In all, these results suggest that RP-832c peptide activity is specific to M2 macrophages.

**Figure 1:**
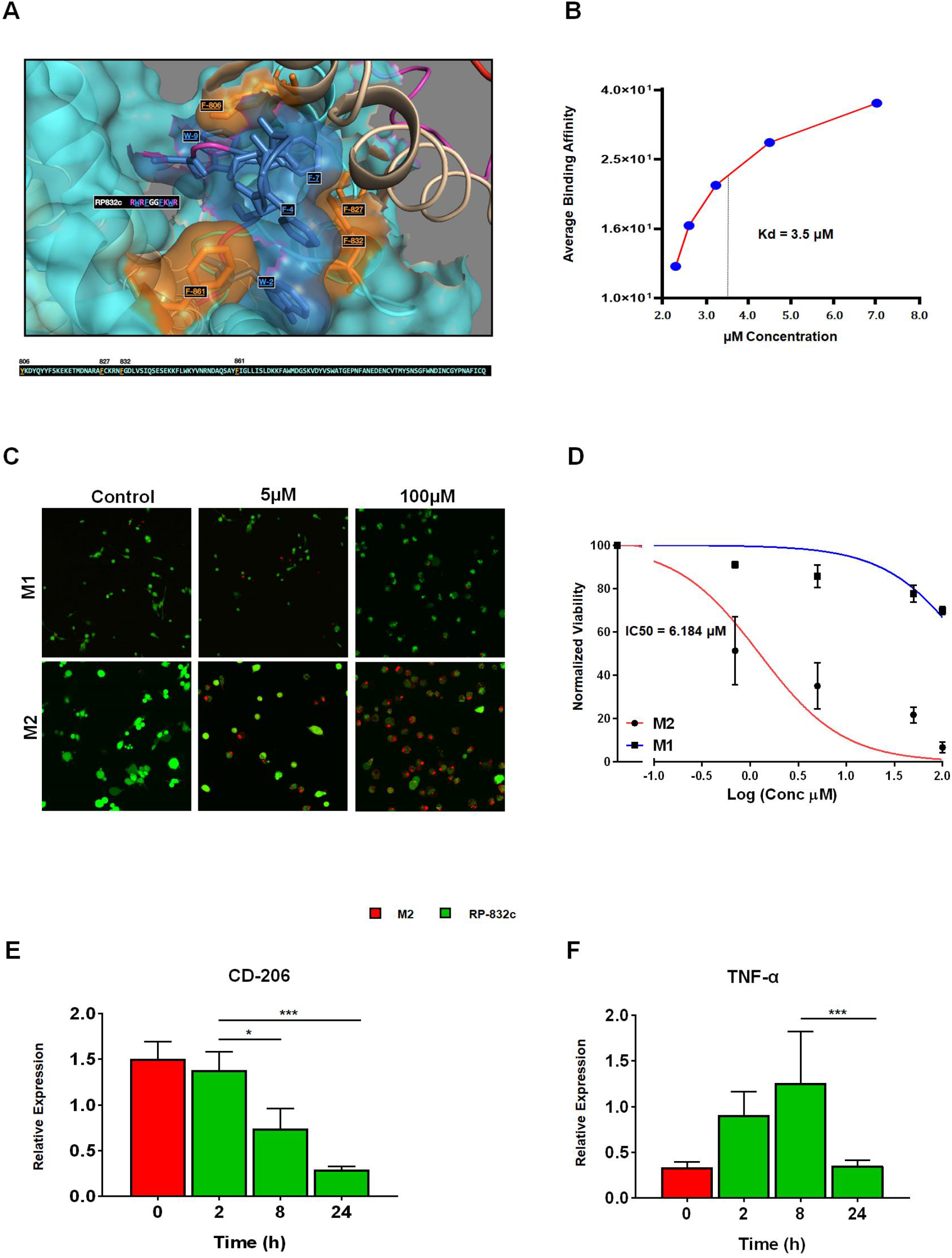
RP-832c specifically targets the CD206 receptor with high affinity. A. In silco docking of RP-832c peptide to the CD206 molecule demonstrates the peptide binds to CD206 with a Binding Energy of −1,349.2 kcal/mol. B. Surface Plasmon Resonance (SPR) spectroscopy analysis for the binding of CD206 protein C. Representative Immunofluorescence images (IF) of Live/Dead assay from BMDM using M1 and M2 macrophage polarization assay. After 48 Hours of treatment with 0μM - 100μM, Dead (red) and Live (green) D. Dose Response Curve of BMDM using M1 and M2 macrophage polarization assay. RP-832c treatment was administered 0μM - 100μM concentration. E. RT-PCR of CD206 gene expression after 0-24 hours of RP-832c treatment on BMDM cells polarized to M2. F. RT-PCR of TNF-α gene expression after 0-24 hours of RP-832c treatment on BMDM cells polarized to M2. All data presented are the means of three independent experiments, performed in triplicate ± S. E. ***P < 0.0001, and ** P<0.001 and *P<.05 is significant.

### RP-832c Prevents Acute Lung Injury in BLM Challenged Mice

We first sought to perform *in silico* validation of CD206 expression at various time points in both bleomycin-treated rat (GSE48455) and mice models (GSE40151). Overall, CD206 expression is significantly upregulated (*p value < 0.0001*) as early as day 2, peaks at day 7 (*p value < 0.0001*), and remains elevated through day 35 (Supplemental Figure 3A, B). We further validated, by immunohistochemistry, that is increased CD206 expression BLM treated mice after 21 days (Supplemental Figure 3C, D). Thus, the BLM model is an appropriate model to determine the role of CD206 lung fibrosis.

To determine the optimal *in vivo* dosage of RP-832c, mice were treated by subcutaneous injection at 5 and 10 mg/kg for a 21 day period following 3 days after a single bolus (2.5U/kg) of BLM. Masson’s Trichrome staining was used to assess lung architecture and collagen deposition. Figure 2A demonstrates that the overall extent of fibrosis is signfiicantly decreased in RP-832c treated mice. Higher resolution images demonstrate that the alveoli of BLM treated mice contained high amounts of fibrotic and collagenous tissue, decreasing the spaces between the alveoli, which are not present in both the 5 and 10mg/kg RP-832c treated mice (Figure 2A), and this further correlates with lower Ashroft scores for both treatment groups (Figure 2D). H&E staining of lung parenchyma lesions from the BLM treated mice tissue sections exhibited variably non-existent lung structure, large fibrotic masses (50% of microscopic field), with only partial lung architecture preserved, and multifocal obliteration of the alveoli by fibrous masses. Moreover, BLM treated mice exhibit eosinophilic, amorphous and slightly vacuolated material [21], as well as thickened alveolar septae with eosinophils, macrophages, lymphocytes and plasma cells similar to those found in ARDS (Figure 2A). In contrast, the RP-832c treated group demonstrated minimal fibrous thickening of alveolar/bronchiolar walls, distended blood vessels and less cellularity. A moderate thickening of alveoli walls without obvious damage was observed in these animals as well (Figure 2A). We further observed a significantly lower overall macrophage count in the RP-832c treated animals compared to BLM challenged or naïve animals (*p-value < 0.001*) (Figure 2E). Interestingly, we observed a similar trend in preventing BLM induced fibrosis for mice treated with RP-832c using intranasal administration. (Supplemental Figure 4). Since 10mg/kg demonstrated the most significant reduction in fibrosis, we used this concentration for the remainder of the experiments.

**Figure 2:**
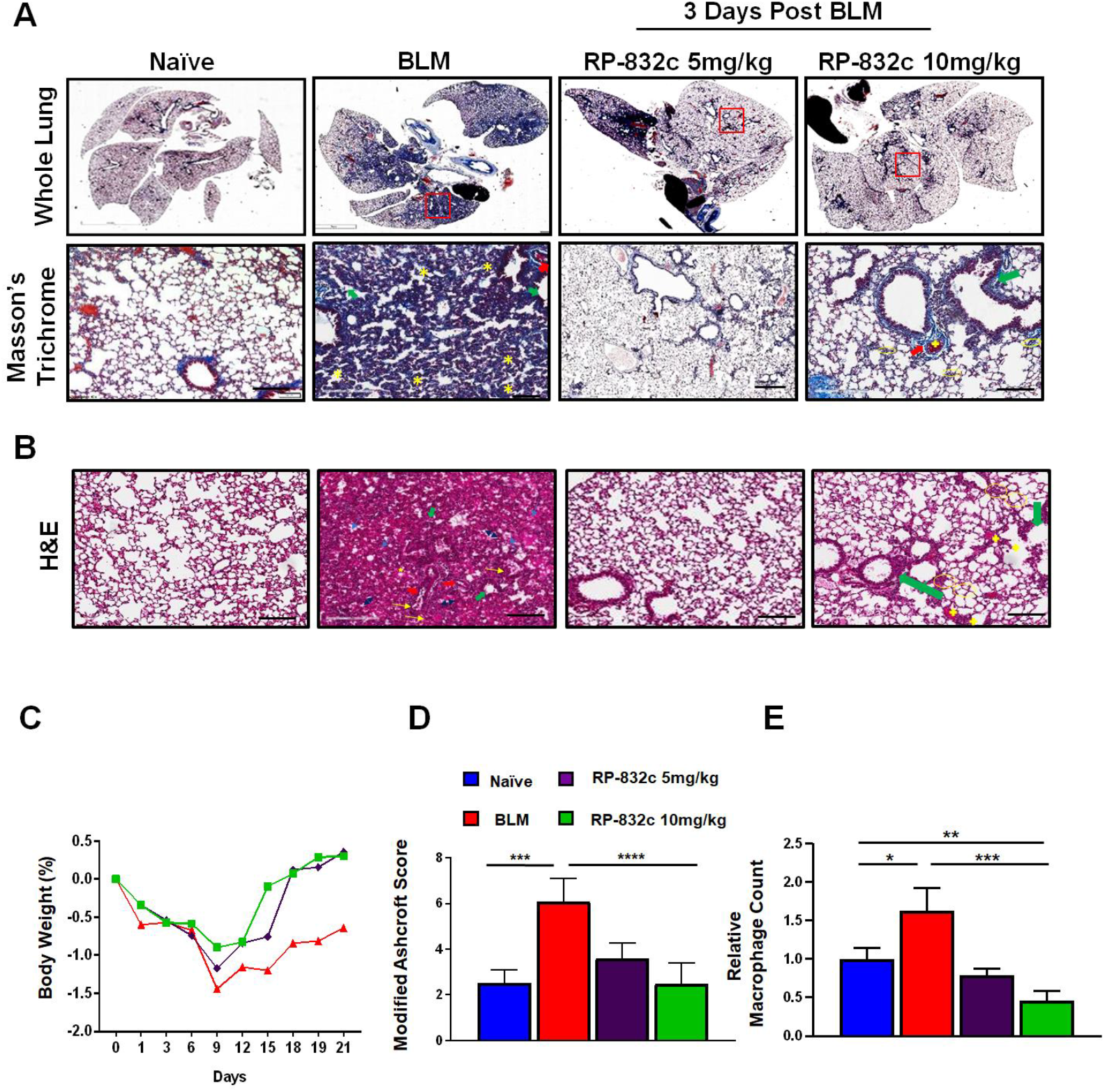
RP-832c peptide prevents fibrosis in an Acute model of lung injury. A. Mice were challenged with a 2.5U/kg body weight dose of BLM for 3 days, after 3 days some of the mice were treated with either 5mg/kg or 10mg/kg RP-832c for an additional 21 days. Upper panel is representative of Masson’s Trichrome staining of whole lung tissue and lower panel is higher resolution images of the area indicated by red box. B. Images are representative of H&E staining of lung tissue sections. Thickening of the bronchiolar wall distended blood vessels with high cellularity. Arrows (Blue) indicate alveolar spaces with edema. Arrows (Yellow) indicate inflammatory cells (lymphocytes, plasma cells, and macrophages); Double arrow ends (Black) indicates some of the distorted and remaining alveolar spaces (air bubbles). Thick arrow(red) indicates perivascular fibrosis. Asterisks show interstitial fibrosis/collagen. Thick arrows (green) indicate parabronchial fibrosis. C. The body weight of the mice was measured over 21 days in each treatment group. D. Lung tissue fibrosis was assessed using the modified Ashcroft scoring system. E. The total number of macrophages was counted microscopically over 10 high fields in the H&E slides. (n=6) mice per treatment group. S. E. ***P < 0.0001, and ** P<0.001 and *P<.05 is significant.

### RP-832c Peptide Reduces CD206 Expression and Fibrosis Markers in Bleomycin Treated Mice

CD206 expression was significantly decreased in RP-832c treated mice compared to BLM challenged mice without treatment (*p-value < 0.001*) (Figure 3A, D). Furthermore, we observed no significant difference in CD170 (a marker for resident alveolar macrophages) expression levels between the RP-832c, BLM or naïve controls (Figure 3A, C). Since, the role of TGF-β1 is well established in promoting fibrosis through inducing a fibroblast to myofibroblast switch, we further measured fibrosis markers TGF-β1 and alpha smooth muscle actin (α-SMA). RP-832c treated lung tissue shows a significant reduction (*p-value < 0.001*) of both α-SMA (Figure 3B, E) and TGF-β1 (Figure 3B) expression compared with BLM treated mice.

**Figure 3:**
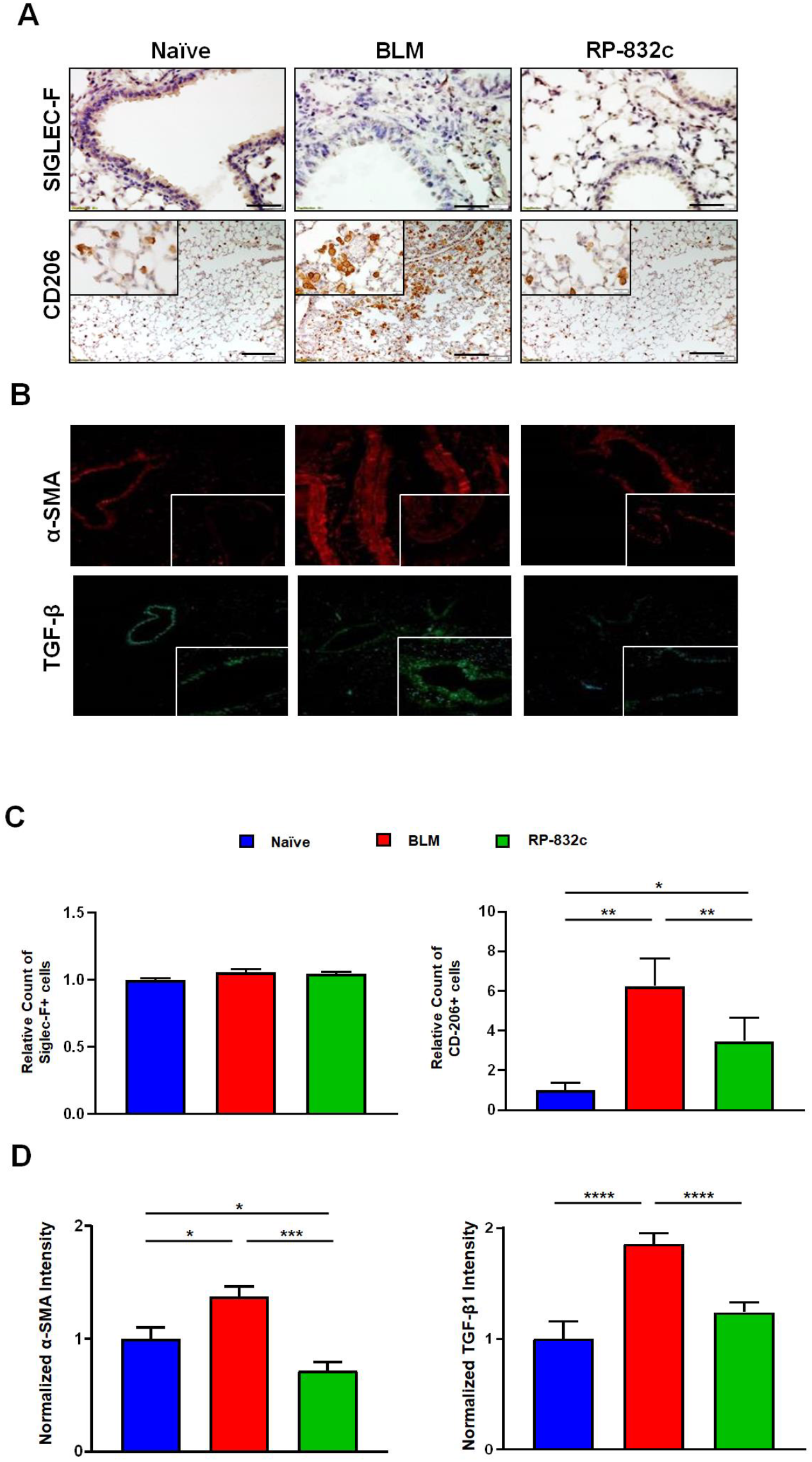
RP-832c peptide significantly decreased CD206+ macrophages and profibrotic markers α-SMA and TGF-β1 without affecting Siglec-F positive alveolar macrophages A. Images are representative of immunohistochemistry staining using Anti-Siglec-F/CD170 or B. Anti-CD206 in naïve, BLM or RP-832c post BLM treated mice. Bar represents 10 μm. C. Lung tissue was stained with Anti-α-SMA (red) and Anti-TGFβ-1 (green) for immunofluorescence and quantified using Metamorph imaging software. The calibration bar represents 20μm. S. E. ***P < 0.0001, and ** P<0.001 and *P<.05 is significant.

To determine the effectiveness of RP-832c on established fibrosis, RP-832c treatment was initiated after 14 days of BLM treatment (Figure 4A). As expected, RP-832c treatment significantly reduced the modified Ashcroft scores compared to BLM only treatment (Figure 4B). Furthermore, we observed significant decreases in CD206 expression by immunofluorescence (Figure 4D). M2 macrophages are responsible for pro-inflammatory and pro-fibrosis cytokines in the lung commonly known as the “cytokine storm” [22, 23] To determine if RP-832c affects these cytokines, we performed RT-PCR on mRNA extracted from naïve, BLM only, and RP-832c treated lungs. Interestingly, we observed significant decreases in TNF-α, IL-10, IL-6, CXCL1, and CXCL2 (Figure 4E). We further observed decreases in M1 macrophages markers IFN-γ, IL-1β and iNOS expression although they did not reach statistical significance (Figure 4E). Concomitantly, fibrosis markers TGF-β1 and MMP-13 were significantly decreased in RP-832c treated mice (Figure 4E). Collectively, our findings demonstrated that a reduction in CD206 positive macrophages correlates with a significant reduction in inflammatory cytokines which in turn reduces myofibroblasts and collagen deposition, which are hallmarks of many types of fibrotic diseases (Figure 6).

**Figure 4:**
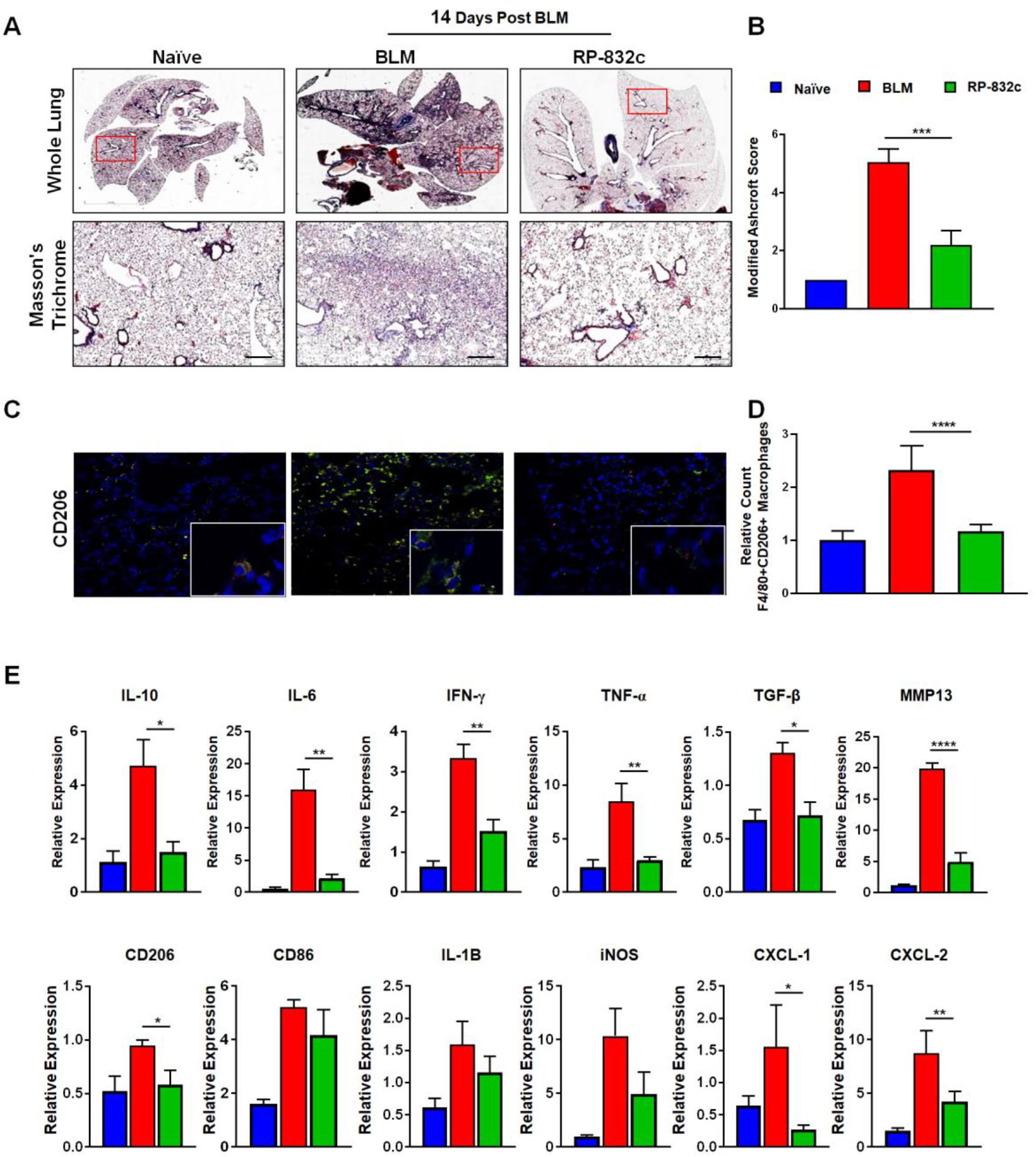
RP-832c treatment blocks the “cytokine storm” in a chronic model of fibrosis. A. At day 0 mice were challenged with a single bolus of 2.5U/kg bwt of BLM, 14 days post BLM challenge mice were treated with 10mg/kg RP-832c for an additional 21 days. Upper panel is representative of Masson’s Trichrome staining of whole lung tissue and lower panel is higher resolution images of the area indicated by red box. B. Lung tissue were scored using modified Ashcroft score. C. Immunofluorescence double staining of lung tissue sections with anti-F4/80 (red) and Anti-CD206 (yellow) D. .CD206 was measured using Metamorph imaging software of 10 separate fields. E) RT-PCR IL-8 functional homologues (CXCL-1, CXCL-2); inflammatory cytokine markers (CD206, CD86, IL-6, IL-10, IFN-γ, TNF-α, IL-1B, iNOS), and fibrosis markers (TGF-β, and MMP-13). S. E. ***P < 0.0001, and ** P<0.001 and *P<.05 is significant.

Pirfenidone and Nintedanib are only two FDA approved drugs for lung fibrosis [24, 25]. Daily treatment of 10 mg/kg of RP-832c treatment was compared to 30mg/kg Pirfenidone or 50mg/kg Nintedanib treatment every three days for 21-day treatment post 14 days of BLM, or in combination (Figure 5A). As a monotherapy RP-832c demonstrated similar Modified Ashcroft score to Pirfenidone treatment (Figure 5B), however in the combination group, there was no significant synergy (Figure 5B). Interestingly, when we compared CD206 expression, RP-823c treatment showed significant decreases in CD206 as expected while the combination of RP-832c with Pirfenidone showed a significant decrease compared to mice in the single therapy groups (Figure 5C). Although these differences were not statistically significant there is a similar trend of synergism in the RP-832c with Pirfenidone combo in decreasing tissue expression of TGF-β1 and α-SMA (data not shown). Interestingly, RP-832c did not synergize with Nintedanib, however RP-832c treatment inhibitor fibrosis to a greater compared Nintedanib only treated mice (Supplemental Figure 5).

**Figure 5.**
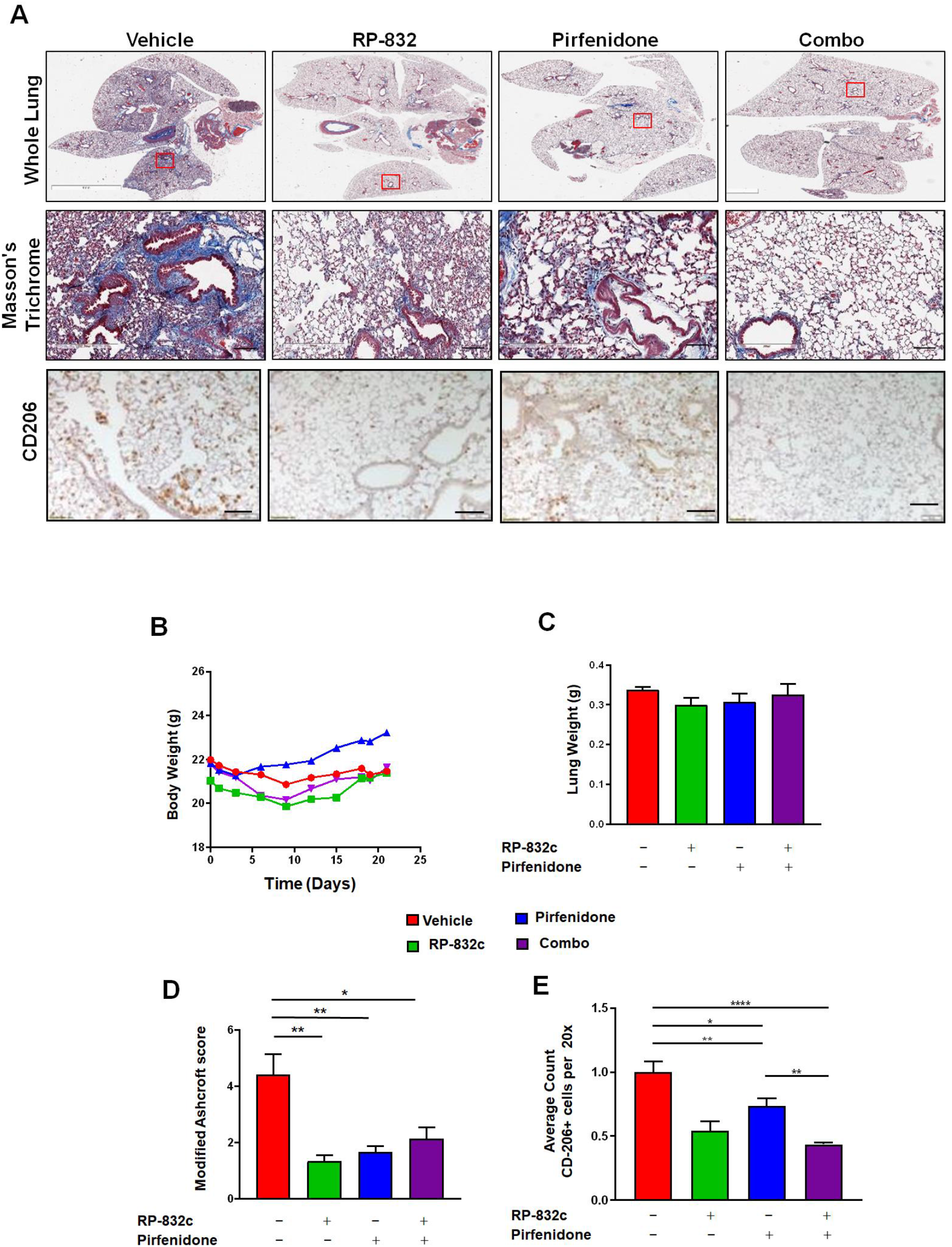
RP-832c peptide shows comparable activity to Pirfenidone in decreasing fibrosis: A. At day 0 mice were challenged with a single bolus of 2.5U/kg bwt of BLM, 14 days post BLM challenge mice were treated daily (QD) with 10mg/kg RP-832c, or every 3 days (QD3) with 30mg/kg Pirfenidone, or a combination for an additional 21 days. Upper panel is representative of Masson’s Trichrome staining of whole lung tissue and below is higher resolution images of the area indicated by red box. The lower panel images represent anti-CD2O6 Immunohistochemical Staining of lung tissue. B. Modified Ashcroft scoring of each treatment group determined after 21 days of treatment. n=6 per treatment group. C. Quantification of lung tissues CD206 expression. The bar represents 10 μm. (n=6) per treatment group. D. The body weights were measured over the course of treatment in each group. E. The lung weights were measured over the course of treatment in each group. S. E. ***P < 0.0001, and ** P<0.001 and *P<.05 is significant.

**Figure 6.**
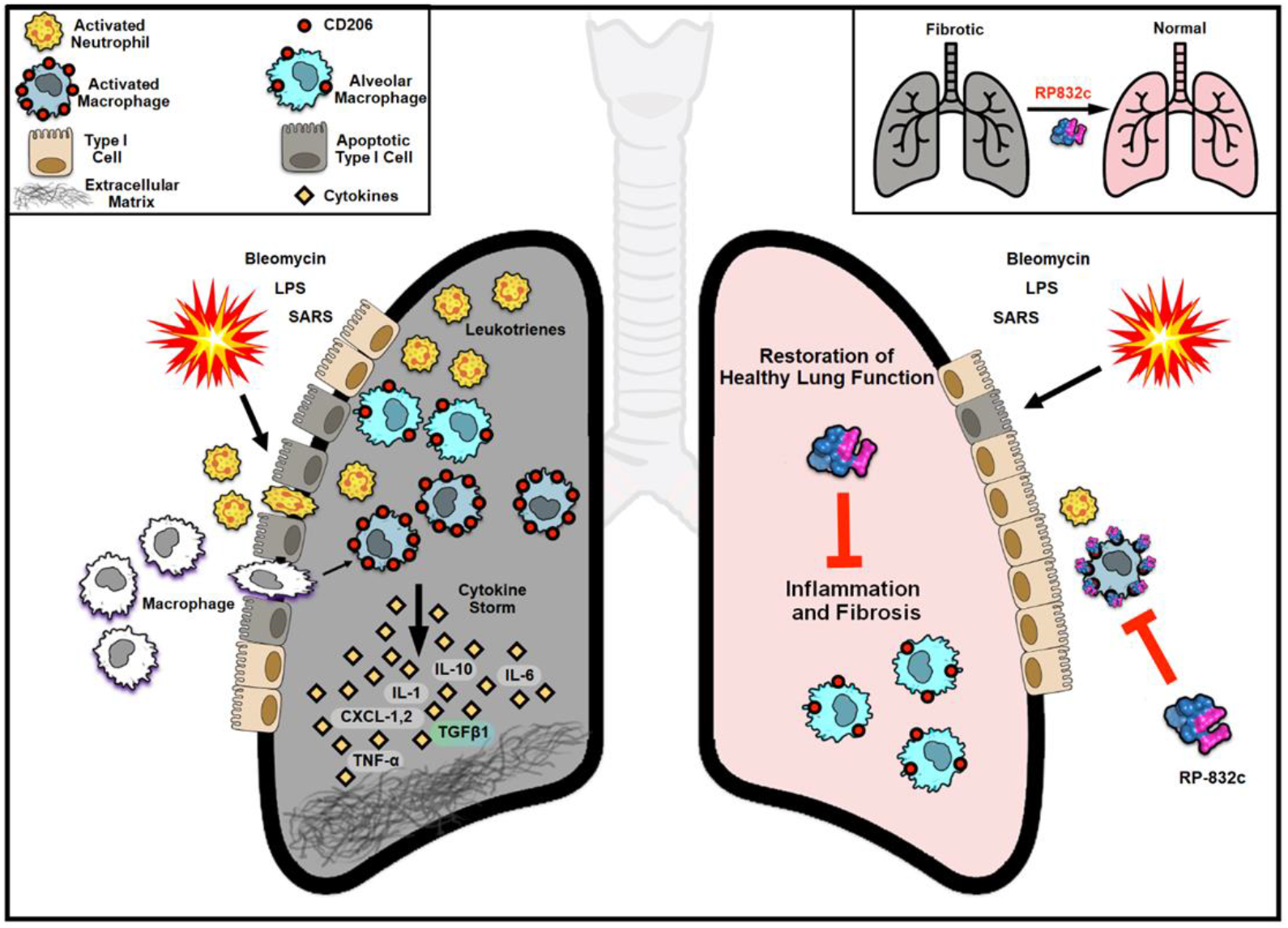
Summary of RP-832c activity on cytokine expression and lung fibrosis. Graphic Summary of lung fibrosis induced by SARS-Cov2, LPS, or Bleomycin (left gray lung panel). During lung injury, circulating monocytes infiltrate the lungs and become activated macrophages. Activated macrophages and leukotrines in turn secrete pro-inflammatory cytokines such as IL10, IL-6, IL-1, CXCL-1,2, TGF-ß, TNF-α which in turn causes inappropriate inflammation that further damages the fibrotic lung. The event is also known as “cytokine storm”. (Right Panel) Targeting CD206 on infiltrating active macrophages with RP-832c suppresses the storm of cytokine, which significantly reduces the inflammatory and fibrotic damages to the lungs and restores normal lung function.

### RP-832c lacks significant toxicity

To determine RP-832c associated toxicity, mice treated with 50 mg/kg was assessed for the body, organ weights, and blood cell compositions. No significant differences were observed (Supplemental Figures 6A & 6B) (Table 1).

**Table 1.**
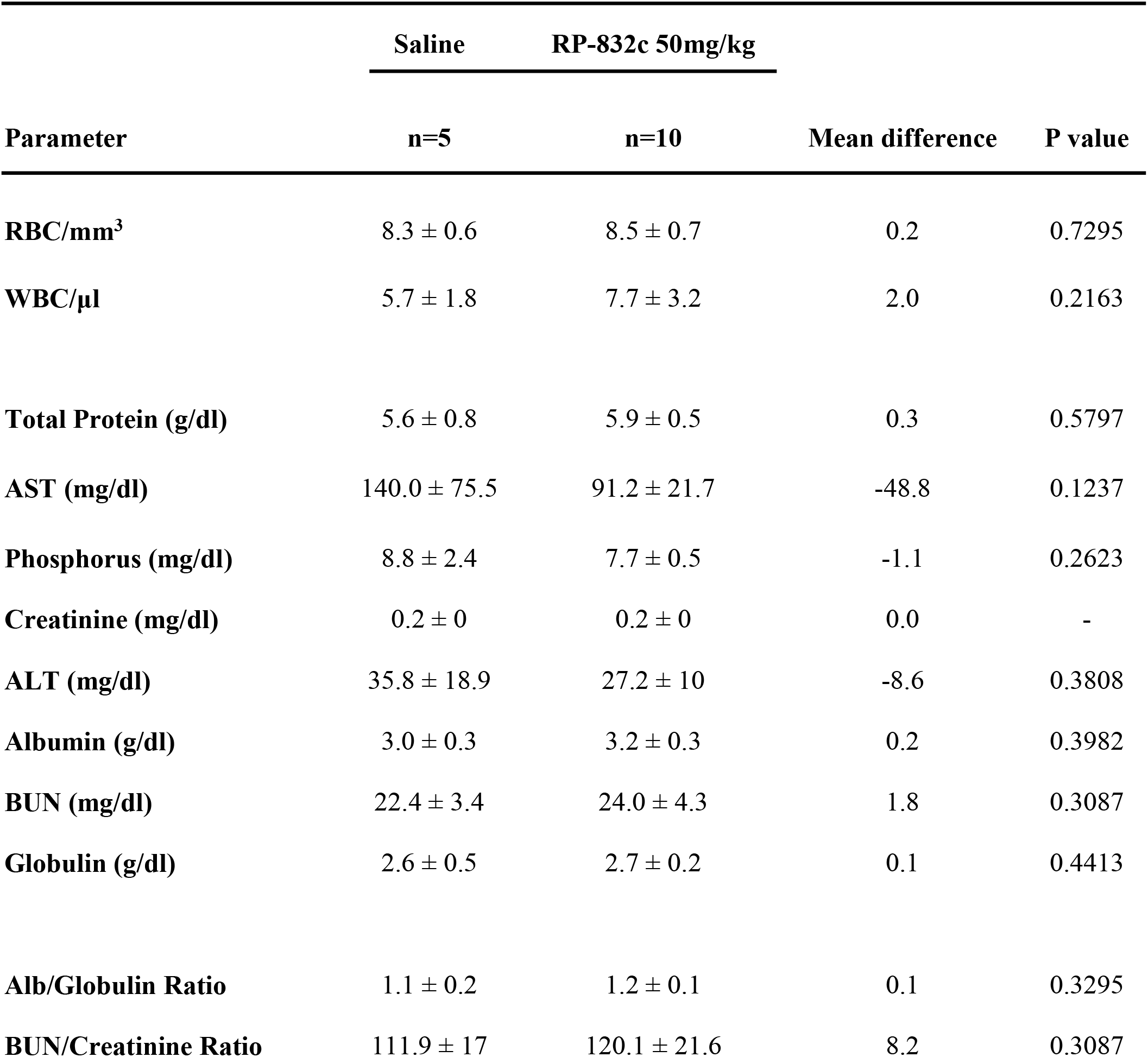
RP-832c peptide does not significantly affect whole blood cell counts.

| **Parameter** | **Saline** | **RP-832c 50mg/kg** | **Mean difference** | **P value** |
| --- | --- | --- | --- | --- |
|  | **n=5** | **n=10** |  |  |
| **RBC/mm<sup>3</sup>** | 8.3 ± 0.6 | 8.5 ± 0.7 | 0.2 | 0.7295 |
| **WBC/μl** | 5.7 ± 1.8 | 7.7 ± 3.2 | 2.0 | 0.2163 |
| **Total Protein (g/dl)** | 5.6 ± 0.8 | 5.9 ± 0.5 | 0.3 | 0.5797 |
| **AST (mg/dl)** | 140.0 ± 75.5 | 91.2 ± 21.7 | -48.8 | 0.1237 |
| **Phosphorus (mg/dl)** | 8.8 ± 2.4 | 7.7 ± 0.5 | -1.1 | 0.2623 |
| **Creatinine (mg/dl)** | 0.2 ± 0 | 0.2 ± 0 | 0.0 | - |
| **ALT (mg/dl)** | 35.8 ± 18.9 | 27.2 ± 10 | -8.6 | 0.3808 |
| **Albumin (g/dl)** | 3.0 ± 0.3 | 3.2 ± 0.3 | 0.2 | 0.3982 |
| **BUN (mg/dl)** | 22.4 ± 3.4 | 24.0 ± 4.3 | 1.8 | 0.3087 |
| **Globulin (g/dl)** | 2.6 ± 0.5 | 2.7 ± 0.2 | 0.1 | 0.4413 |
| **Alb/Globulin Ratio** | 1.1 ± 0.2 | 1.2 ± 0.1 | 0.1 | 0.3295 |
| **BUN/Creatinine Ratio** | 111.9 ± 17 | 120.1 ± 21.6 | 8.2 | 0.3087 |

## Discussion

Macrophages demonstrate remarkable plasticity and can acquire phenotypes which both drive and resolve fibroproliferative responses to injury. One key role of M1 macrophages is to initiate the host defense against pathogens by generating reactive nitric oxide (NO) via inducible nitric oxide synthase (iNOS) and releasing proinflammatory cytokines and chemokines such as IL-1β, IL-12, IL-23, CCL2 and TNF-α, while M2 macrophages contribute to the pathogenesis of fibrosis, by secreting anti-inflammatory and profibrotic growth factors such as TGF-β1 [26]. To modulate these M2 macrophages, we previously reported the development of a first in class HDP derived peptides, RP-182 and RP-832c which specifically target the CD206 receptor on M2-macrophages [18]. These peptides are designed to specifically target the CRD5 domain of the CD206 receptor on M2 macrophages, which induces rapid folding of the CD206 receptor, as well as a decrease of CD206 expression and increase in M1 Marker CD86 expression and apoptosis of CD206 positive population of cells[18]. This current report specifically focuses on RP-832c, selected based on its enhanced stability and increased half-life compared to RP-182 (data not shown). RP-832c treatment induced a significant decrease in CD206 expression levels in both the 3-day and 14-day bleomycin-challenged mice, as well as significantly decreased fibrosis scores, suggesting that targeting elevated CD206 positive M2 macrophages contributes to decreases in lung fibrosis.

SARS-CoV-2 infected individuals that experience ARDS experience a surge in IL-6, IL-10 and TNF-α levels during illness which declines during recovery [27]. In our 14-day BLM model of ARDS associated fibrosis, RP-832c significantly decreases the mRNA expression levels of multiple cytokines including TNF-α, IL-6, IL-10, and IFN-γ in the lung. Since TNF-α induces neutrophil migration, we further observed significant decreases in CXCL1, and CXCL2 in RP-832c treated mice. While each of these markers appears to return with time to naïve mice levels, M1 markers iNOS, CD86, and IL-1β remained relatively higher than in naïve mice. This is not surprising since we previously reported that the RP-182 induces rapid repolarization to the M1 phenotype, and trigger apoptosis in a subpopulation of CD206 positive cells [18]. This increase in M1 macrophages and decrease in CD206 positive M2 macrophages, with either RP peptide treatment or in CD206 KO mice, is associated with an increase of CD8+T cells,[18]. Severely affected patients, that require ICU admission, develop a similar storm of cytokines, which blocks CD4+ and CD8+ T cell infiltration[27]. Although this report focuses on the role of macrophages in ARDS, it is plausible that targeting CD206 positive macrophages with HDPs, will have a similar effect in increasing CD4+ and CD8+ T cell count in the lung, as we observed in cancer models. The fact that the overall lung architecture of RP-832c treated mice is similar to naïve lung, further highlights the resolving effect of RP-832c peptide administration.

A critical factor in macrophage associated lung fibrosis is the origin of M2 macrophages. In the normal injury-repair response, macrophages readily acquire an M2-phenotype which promotes fibroproliferation [28]. These M2 polarized macrophages promote collagen synthesis and deposition as well as remodeling of the extracellular matrix that contributes to the buildup of fibrotic tissue [6]. It is well characterized that CD206 is expressed on both tissue resident alveolar macrophages as well as infiltrating monocyte derived alveolar macrophages [29]. Our findings demonstrate that RP-832c did not influence the expression of Siglec-F/CD170, a marker for resident alveolar macrophages in the lung, suggesting that RP-832c targets CD206 positive macrophages derived from infiltrating monocytes, which have been suggested to contribute to fibrosis. In support of this are two lines of evidence. First, Misharin et al. [30] demonstrated in BLM mouse models of lung fibrosis, that the deletion of monocyte derived alveolar macrophages after their recruitment to the lung markedly attenuated the severity of fibrosis, whereas the deletion of tissue resident alveolar macrophages had no effect on fibrosis severity [30]. Second, depletion of circulating monocytes using CCR2–/– mice or the administration of liposomal clodronate reduces fibrosis severity, implicating monocyte-derived cells in the development of fibrosis [31]. Furthermore, we did not observe any significant effect on the proliferation of multiple fibroblast cell lines cultured in vitro. Thus, it appears that RP-832c specifically affects monocyte-derived M2 polarized macrophages and does not reduce the protective resident alveolar macrophages.

Since infiltrating macrophages are critically responsible for contributing to fibrosis as seen in IPF and SARS-CoV-2 patients with ARDS, several reports have demonstrated that production of profibrotic cytokines like TGF-β1 and IL-4 polarize monocytes and M1 cells towards the M2-like phenotype upregulating CD206 expression [32, 33], secreting cytokines that promote fibroblast to myofibroblast transition [34]. Similarly, injection of anti-TGF-β1 antibody in BLM treated mice resulted in a 40% reduction in collagen accumulation [35]. Interestingly, RP-832c treatment of mice resulted in a decrease in the thickness of the alveolar/bronchiolar wall compared to BLM alone. RP-832c also reduced collagen deposition and fibrosis by 30%, comparable with the effect of directly blocking TGF-β1, one of the most important molecules in promoting fibrosis. Multiple reports demonstrate that the CD206 receptor is critical for ECM remodeling by macrophages and fibroblasts, through upregulation of MMPs, namely 7, 11, and 13 which promote collagen turn over [36, 37]. Indeed, we observed significant decreases in MMP-13, as well as decreases in TGF-β1, and α-SMA tissue expression which correlated with reduction in the myofibroblast cells that are responsible for excessive collagen deposition and tissue remodeling of pulmonary fibrosis.

Nintedanib and Pirfenidone are the only are approved compounds for lung fibrosis, and there are several case reports of their usage on ARDS patients [38]. RP-832c demonstrates a significant decrease in fibrosis compared to Nintedanib with comparable efficacy Pirfenidone in BLM treated mice. Although the mechanism of action of Pirfenidone is not known, we observed decreased CD206 expression in both RP-832c and Pirfenidone, with a trend toward significance in the combination treatments. Interestingly, a recent report noted that while Nintedanib significantly reduced BAL lymphocytes and neutrophils, it did not have an effect on macrophages [25]. Therefore, the significant effect of RP-832c on fibrosis is likely to be through directly targeting M2 macrophages. Although, TGF-β antibodies have been proposed as an anti-fibrosis treatment, they have not fared well in the clinic. Targeting CD206 expression on M2 macrophages with RP-832c can serve as inhibition of the cellular source of TGF-β, with similar effects as targeting TGF-β directly.

In summary, HDPs offer significant anti-fibrotic activity with the advantage of no observed toxicity at 5 times therapeutic inhibitory dosages. Immunotherapies are now being proposed as treatment options for IPF patients. RP-832c appears to fit in this class of immunotherapies and could serve as a potential option for treating pulmonary fibrotic illnesses such as ARDS in SARS-CoV-2/COVID-19 patients.

## Acknowledgments

The authors would like to thank all those who contributed technically and financially to the completion of this project. The authors would like to thank Rosio Monroy, Maria Monroy and Adriana Ledesma at Murigenics, Inc for their help with intratracheal bleomycin administration of the mice.

## Author contributions

JMJ designed the peptides. GM, and HL designed the studies, and animal studies in were performed SN, CK, and AG, performed the staining and analysis. AS conducted in silico binding studies. In vitro experiments were performed by AG, AS, HL, AG performed bioinformatic analysis of GEO datasets and RNA sequencing gene expression data was conducted by AG, and histopathology by AG and BK. BA performed the tissue scoring and pathological analysis. CY, HL and GM provided funding, JMJ, UR, RS, AG, GM, HL and CY wrote the manuscript.

