## Supplemental Figure 1 for "A Novel CD206 Targeting Peptide Inhibits Bleomycin Induced Pulmonary Fibrosis in Mice"

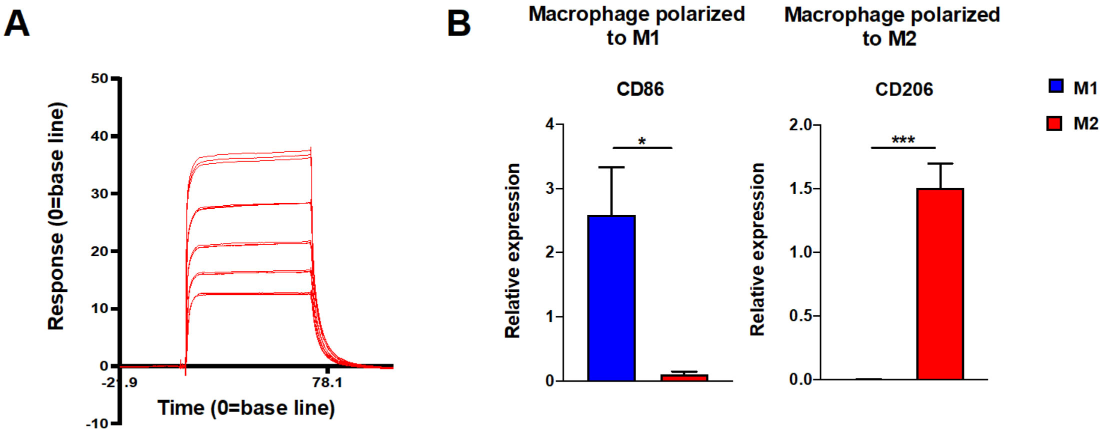


**Supplemental Figure 1.** Dose Response Curve of BMDM and Validation of in vitro BMDM polarization assay. A) SPR (Surface plasmon Resonance) Dose response for the inhibition of CD206 receptor with different concentrations (0µM - 100µM concentration.) of RP-832c B) Validation of in vitro BMDM using C57BL/6 mice polarization assay. Bone marrow derived macrophages polarized to M1 macrophages with INF-γ showed increased CD86 expression while those polarized to an M2 phenotype using IL4 stimulation showed increased CD206 expression.
