## Supplemental Figure 2 for "A Novel CD206 Targeting Peptide Inhibits Bleomycin Induced Pulmonary Fibrosis in Mice"

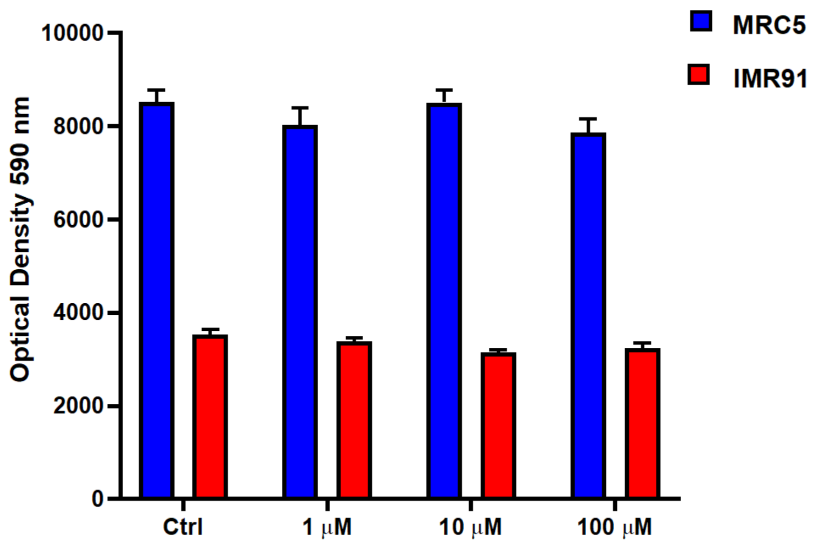


**Supplemental Figure 2.** RP-832c shows no cytotoxicity against lung fibroblasts: RP-832c did not show any cytotoxicity against lung fibroblast cell lines MRC5 and IMR91. All data presented are the means of three independent experiments, performed in triplicate ± S. E. ***P < 0.0001, and ** P<0.001 and *P<.05 is significant.
