## Supplemental Figure 3 for "A Novel CD206 Targeting Peptide Inhibits Bleomycin Induced Pulmonary Fibrosis in Mice"

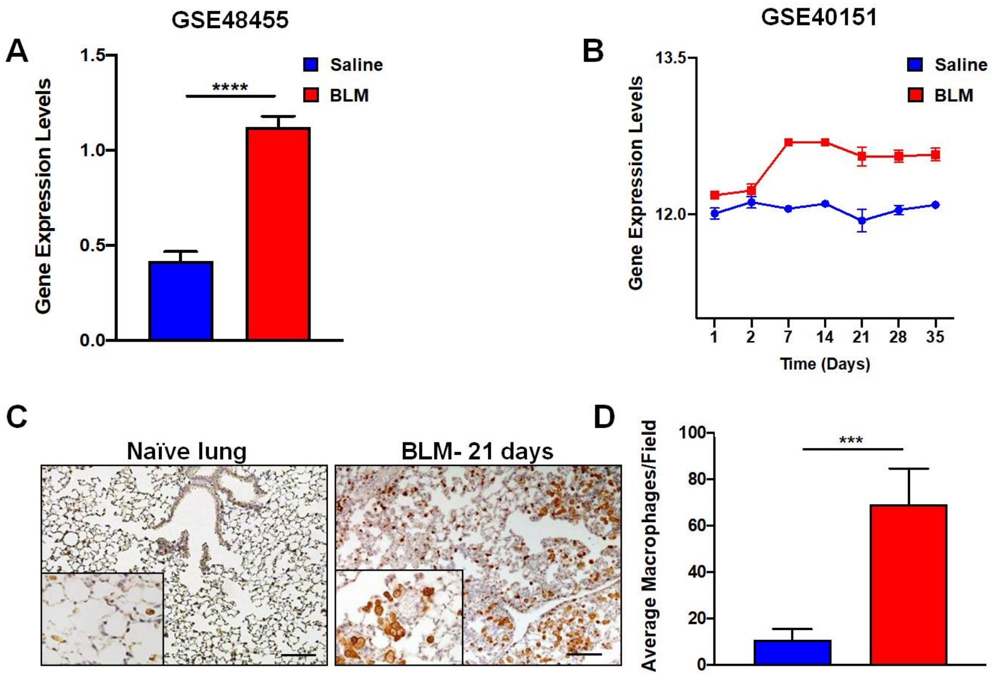


**Supplemental Figure 3.** CD206 expression is increased by BLM treatment. A. Secondary analysis of (GSE48455) data set with 72 mice for CD206 BLM treated compared to normal saline) administered mice. B. Secondary analysis of (GSE40151) of 111 mice were treated with BLM or normal saline for 35 days and mRNA levels CD206 expression were analyzed. C. Images represent anti-CD206 staining on BLM treated mice compared to control. D. CD260 was measured by immunohistochemistry after 21 days of BLM treatment compared to naïve control in n=3 mice ***P < 0.0001, and ** P<0.001 and *P<.05 is significant.
