## Supplemental Figure 4 for "A Novel CD206 Targeting Peptide Inhibits Bleomycin Induced Pulmonary Fibrosis in Mice"

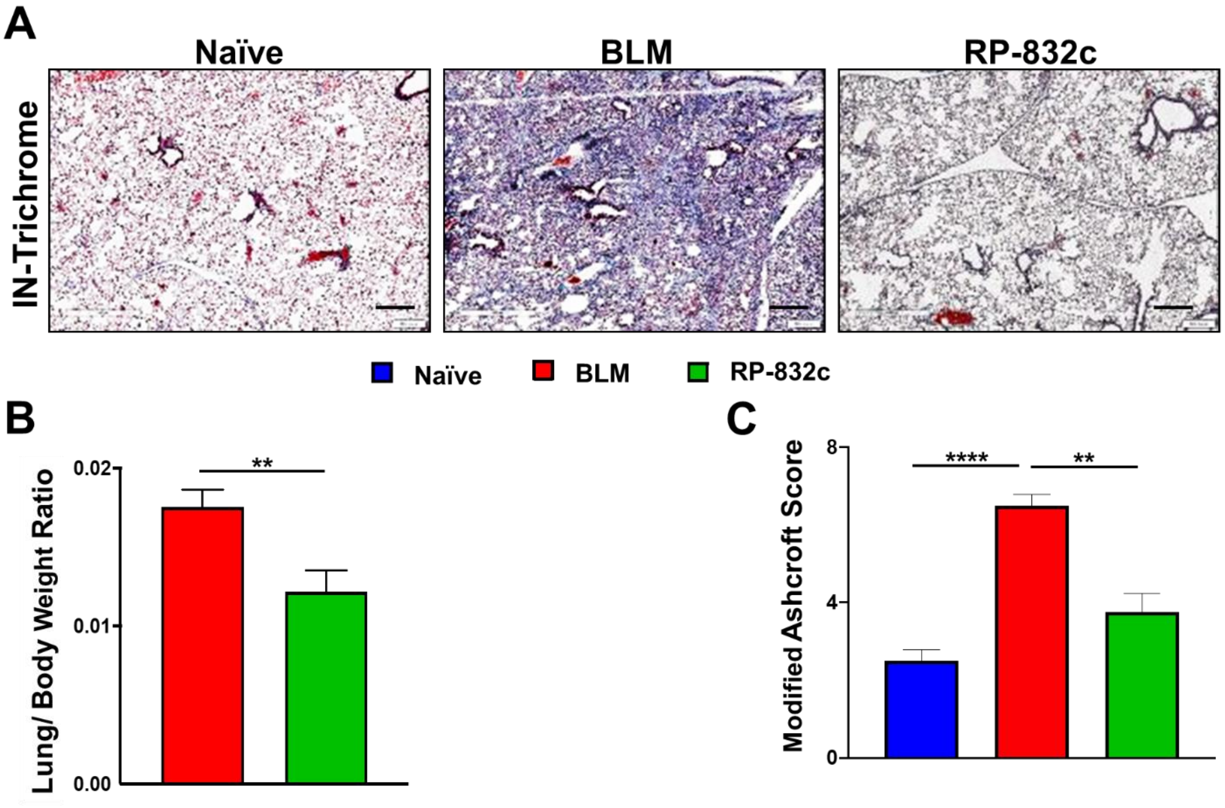


**Supplemental Figure 4.** Intranasal administration of RP-832c peptide shows similar activity to the subcutaneous administration route. A. RP-832c peptide was administered intranasally at 2.5µg/mouse QD through day 18. B. The lung to body weight ratio of lung tissues from the different treatment groups was compared. C. Images are representative of Masson's trichrome staining of Intranasally administered RP-832c. D. Modified Ashcroft score of Intranasally administrated RP-832c compared to vehicle control.
