## Supplemental Figure 5 for "A Novel CD206 Targeting Peptide Inhibits Bleomycin Induced Pulmonary Fibrosis in Mice"

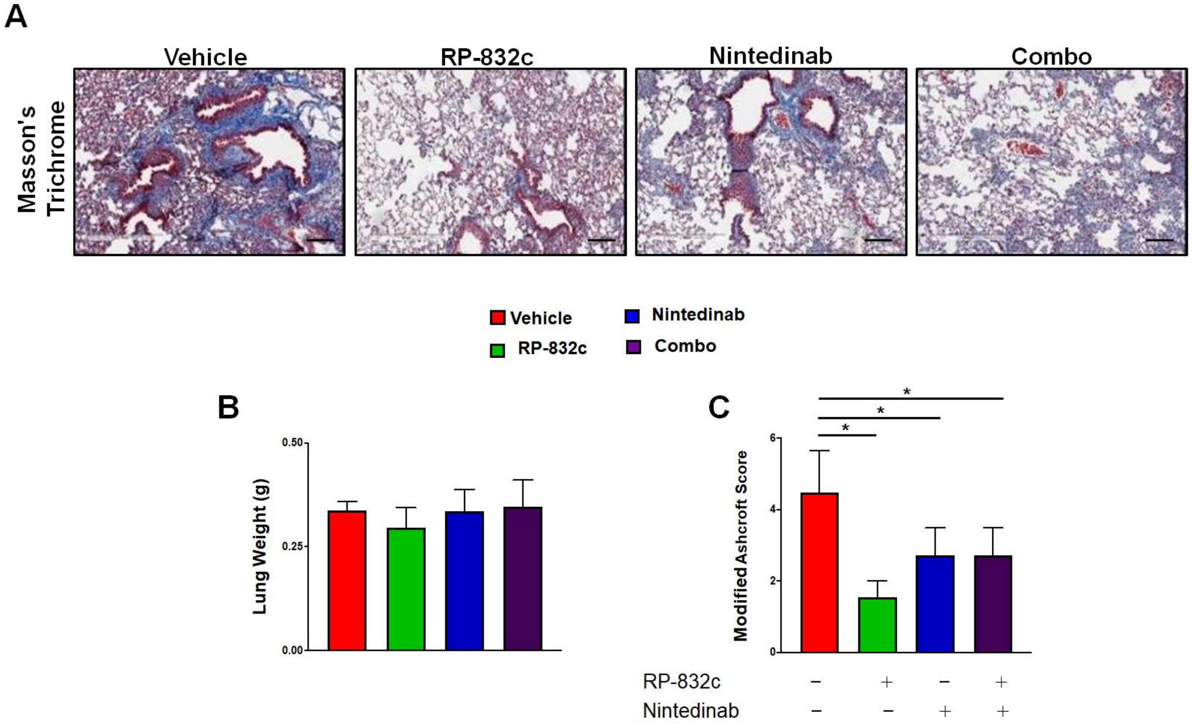


**Supplemental Figure 5.** RP-832c peptide decreased fibrosis to a greater extent compared to Nintedanib. At day 0 mice were challenged with a single bolus of 2.5U/kg bwt of BLM, starting 14 days post BLM challenge mice were treated daily (QD) with 10mg/kg RP-832c, or every 3 days (QD3) with 50mg/kg Nintedanib, or in combination for an additional 21 days. Masson's Trichrome staining of lung tissue B. The lung weights were measured over the course of treatment in each group. C. Modified Ashcroft scoring of each treatment group was determined after 21 days of treatment. n=6 per treatment group. S. E. ***P < 0.0001, and ** P<0.001 and *P<.05 is significant.
