## Supplemental Figure 6 for "A Novel CD206 Targeting Peptide Inhibits Bleomycin Induced Pulmonary Fibrosis in Mice"

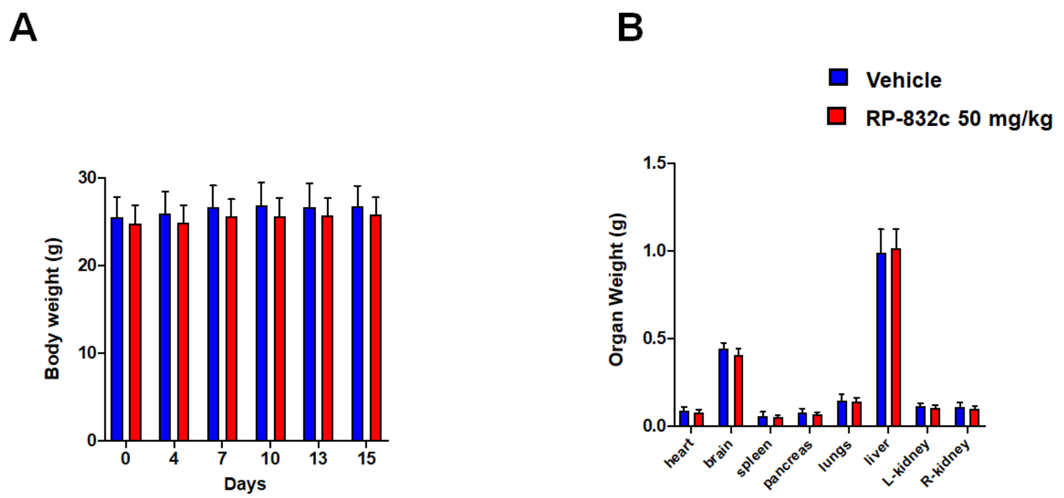


**Supplemental Figure 6:** RP-832c peptide lacks toxicity A) Animal toxicity study conducted using CD1 mice showed that 50mg/kg RP-832c treatment did not have any significant effect on body weight compared to normal saline treated mice. B) Additionally, 50 mg/kg of RP-832c also does not have any significant effect on organ weight compared to saline treated mice.
